# Cas9-expressing HC-04 hepatocytes facilitate CRISPR-based analysis of *Plasmodium falciparum* sporozoite-host interactions

**DOI:** 10.1101/2025.08.19.670452

**Authors:** Eva Hesping, Lisa H. Verzier, Marcel Doerflinger, Marco J. Herold, Justin A. Boddey

## Abstract

Sporozoites of *Plasmodium falciparum*, the deadliest malaria parasite, are transmitted into the skin by infected mosquitoes and migrate to the liver to initiate infection. There, they invade hepatocytes and develop into exoerythrocytic merozoites that, eventually, enter the bloodstream and invade erythrocytes, leading to malaria. The parasite journey involves cell traversal, where sporozoites transiently enter and exit host cells beginning in the skin, lysing membranes to move deeper into tissue and evade immune cell destruction. After reaching the liver and traversing several hepatocytes, sporozoites productively invade a final hepatocyte to establish liver-stage infection. The molecular mechanisms underlying traversal, invasion, and intracellular development remain incompletely understood, particularly with respect to host determinants. To address this, we engineered human HC-04 hepatocytes, the only known cell line supporting *P. falciparum* liver-stage development, to express Cas9-mCherry, enabling CRISPR-based functional genomics studies. We validated Cas9 activity and demonstrated successful guide-RNA-directed gene disruption via non-homologous end joining in HC-04 Cas9+ (clone 2B3) cells. Optimized traversal and invasion assays with HC-04 2B3 cells led to a robust cytometric assay suitable for screening human genes involved in *P. falciparum* infection. As proof-of-concept, we performed a small screen involving disruption of 10 human genes previously implicated in infection by bacterial and viral pathogens, confirming utility of this platform. While no new host factors were identified for malaria parasites in this initial study, we have developed a tractable system for genome-wide CRISPR screens to uncover novel hepatocyte biology and host determinants of infection by liver-tropic pathogens.

## Introduction

Among the *Plasmodium* species responsible for human malaria, *P. falciparum* is the most prevalent, accounting for over 95% of cases and deaths globally (1). Transmitted by the bite of an infected *Anopheles* female mosquito, *P. falciparum* sporozoites are deposited into human skin during blood feeding. To establish infection, sporozoites must migrate through the dermis and enter a blood vessel for transport via the bloodstream to the liver sinusoids. There, they migrate across the endothelium into the liver parenchyma, traverse hepatocytes and then invade a final hepatocyte to continue their lifecycle (2–4). *Plasmodium* sporozoites have evolved to interact with their host in multiple ways. First, through gliding motility that is characterised by start-and-stop movements due to successive attachment and cleavages of surface adhesins that enable migration through host tissues (3–5). They have also evolved a remarkable mechanism called cell traversal, which allows them to cross physical barriers and avoid destruction following uptake by immune cells. Cell traversal refers to the sporozoites’ ability to pass through cells, entering and rapidly exiting them through the perforation and destabilization of host cell membranes that are rapidly repaired following the sporozoite breach (6–8). Cell traversal is critical to infect mammalian hosts, as parasites deficient in a group of proteins related to the ‘membrane attack complex/perforin (MACPF)’ family are unable to efficiently traverse cells or infect the liver, even when delivered intravenously to bypass the dermal barriers (9–13). Interestingly, even after reaching the liver, sporozoites traverse multiple hepatocytes before finally committing to productive invasion of a single hepatocyte. Parasites establish and develop within a parasitophorous vacuole membrane (PVM) over the next 7 days before egressing and releasing merozoites, the form that infects erythrocytes (6,14,15). In contrast to productive invasion, traversing parasites briefly occupy a compartment called a transient vacuole (TV). Unlike the parasitophorous vacuole (PV) formed during productive invasion, the TV is marked by the presence of F-actin and phosphatidylinositol-3-phosphate (PI3P) and is rapidly lysed, whereas the PV lacks both proteins and remains intact (16,17) unless attacked by cell-autonomous innate responses that clear the infected cell (18).

Several pore-forming and MACPF-like proteins have been identified and are essential to the process of cell traversal. Sporozoite microneme protein essential for cell traversal (SPECT) and perforin-like protein 1 (PLP1, also called SPECT2) are pore-forming proteins that play key roles in cell traversal. Initially identified in rodent malaria models (9–12,19), both were later confirmed to be essential for *P. falciparum* cell traversal and sporozoite infectivity of humanized chimeric liver mice (12). PLP1 has been shown to be essential during parasite egress from the TV, underlining its role in traversal (16). Cell traversal protein for ookinetes and sporozoites (CelTOS) is another crucial factor for parasite motility and host membrane disruption and is involved in traversal by both ookinetes at the mosquito midgut and sporozoites in the mammalian host (20,21). Biochemical studies revealed that CelTOS forms ∼50 nm pores in diameter through preferential binding to phosphatidic acid, a lipid component enriched on the inner leaflet of host cell membranes. Its role in parasite exit from host cells has, however, yet to be conclusively demonstrated (22). How these pore-forming proteins coordinate to disrupt host cell membranes, whether acting in concert or in sequence, remains unresolved. In addition, AMA1 and MAEBL, more commonly associated with invasion of mosquito salivary glands and erythrocytes have also been implicated in cell traversal and productive invasion of hepatocytes by *P. falciparum* sporozoites (23–25). Despite the importance of cell traversal in establishing infection in the mammalian host, little is known about the host molecular mechanisms underlying this process. The overlap between parasite proteins involved in cell traversal and productive invasion suggests that these pathways may share common host-parasite interactions. Supporting this, several studies that did not differentiate between traversal and invasion have identified host cell processes, such as exocytosis and actin polymerization, as essential for sporozoite entry into hepatocytes by *Plasmodium* species that infect rodents (26,27). Although the molecular determinants differentiating cell traversal from invasion remain incompletely defined, these findings imply convergence in the cellular machinery that may be exploited during both processes.

Host factors implicated in *Plasmodium* invasion differ across parasite and host species and model systems. CD81, the first host protein identified as essential for *P. falciparum* invasion of primary human hepatocytes (28,29) is also important for *P. yoelii* sporozoite invasion and rhoptry discharge in rodent models (30,31). Other host receptors implicated in hepatocyte invasion or development were identified through genetic screening. Host factors involved in invasion and development of sporozoites were identified by short interfering RNA (siRNA) screens, identifying scavenger receptor class B type I (SR-BI) as important for *P. berghei*, and later by the human malaria parasite *P. vivax* (32–35), and PKCζ, involved in rodent malaria parasite growth (36). Another approach involved comparing hepatocytes that were permissive, or not, to sporozoite infection to identify host proteins required for infection (35,37–39). More recently, CRISPR approaches that offer robust and long-lasting gene disruption have been used to identify host factors for invasion and intracellular development of infectious pathogens, including apicomplexans such as *P. yoelii* (36,40–43); however, CRISPR screens have not yet been reported with *P. falciparum* sporozoites. In *P. yoelii*, a CRISPR screen identified centromere protein J (CENPJ) as a regulator of rodent malaria parasite development in the liver (44). This screen was performed in HepG2 human hepatocytes, which are useful for *Plasmodium* species that infect rodents but are not permissive to productive infection by *P. falciparum* sporozoites (45).

Human HC-04 cells remain the only hepatocyte cell line capable of supporting *P. falciparum* liver-stage development, albeit with invasion and intracellular development efficiencies that are imperfect (13,46–49). While HC-04 cells do not express appreciable levels of CD81 (47), *P. falciparum* invades and develops into exoerythrocytic forms within these cells (46). Additionally, anti-CD81 antibodies do not block invasion in this model, suggesting the parasite may exploit CD81-independent entry pathways. Despite these limitations, HC-04 cells have been commonly used to investigate *P. falciparum* pre-erythrocytic biology (13,46–49), and although the influence of experimental variables on *P. falciparum* HC-04 interactions has been explored (37,47), many aspects remain incompletely understood. We therefore engineered Cas9-expressing HC-04 cells and validated clone 2B3 for CRISPR-based functional genomics studies in the context of *P. falciparum* liver-stage infection. Cas9+ HC-04 2B3 are a valuable cell line tool for genome-wide and arrayed CRISPR screens to uncover host determinants of liver-stage infection by human malaria parasites, including *P. falciparum* and *P. vivax*.

## Results

### Generation of Cas9+ HC-04 hepatocyte clones with endonuclease activity

Given their prior use to investigate cell traversal and invasion by sporozoites (12,47–50), HC-04 hepatocytes were selected to enable functional studies of *P. falciparum*-hepatocyte interactions by genome editing. To this end, Cas9-expressing HC-04 cell line clones were generated by transduction of wild-type (WT) HC-04 cells with a lentiviral construct encoding Cas9 fused to the fluorescent protein mCherry separated by a 2A skip peptide (Cas9-mCherry). Successfully transduced single-cell clones were isolated by fluorescence-activated cell sorting (FACS) and expanded in independent wells. Three Cas9-mCherry+ clones, 1C3, 1C6, and 2B3, were identified by mCherry+ signal and selected for further characterisation (**Figure 1a**).

**Figure 1:**
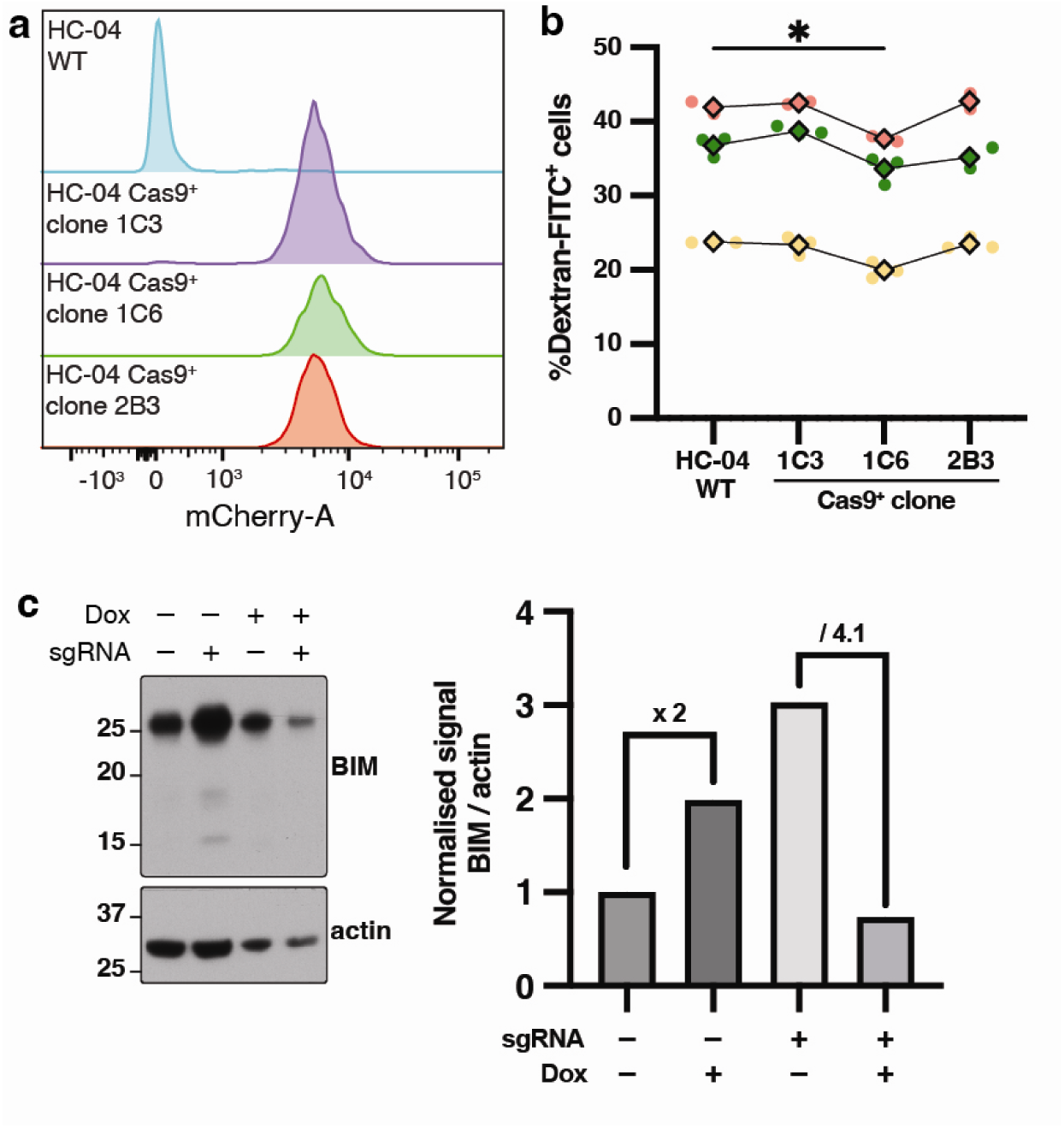
Generation of Cas9-mCherry+ HC-04 clones. **(a)** Cas9-mCherry+ HC-04 clones 1C3, 1C6 and 2B3 express mCherry+ as shown by flow cytometry. (**b**) Number of dextran+ (traversed) HC-04 cells. Only a modest reduction for clone 1C6 was observed (p= 0.0135, Dunnett multiple comparisons test). Colours represent biological replicates with both technical replicates (round points) and their average (diamond) shown. The multiplicity of infection (MOI) was 1, except in yellow MOI0.5 was used. **c**. Immunoblots show expression of BIM and actin in HC-04 2B3 transduced with sgRNA targeting *BIM* under expression of a doxycycline-inducible promoter (left). Densitometry of anti-human BIM and anti-human actin antibodies is shown (right) and the fold-change indicated numerically (above).

To ensure that Cas9-mCherry expression did not affect *P. falciparum* sporozoite cell traversal, Cas9-mCherry+ HC-04 clones were assessed for traversal by sporozoites and compared to WT parental HC-04 cells by measuring their permissiveness to the uptake of dextran-FITC that is otherwise impermeable to HC-04 cells (12,13). The rate of cell traversal for WT HC-04 and Cas9+ clones 1C3 and 2B3 was similar, while sporozoites traversed clone 1C6 at a modestly reduced rate (**Figure 1b**, *p*=0.0135). Clone 2B3 was selected for further characterization.

To confirm that Cas9 was functional when expressed in HC-04 cells, the endonuclease activity of clone 2B3 (hereafter called HC-04 2B3) was investigated following transduction of a doxycycline-inducible single guide RNA (sgRNA) targeting the human *BIM* gene or a mock transduction lacking a sgRNA as a control (51). The *BIM* gene was selected as reagents were readily available to study it, and sgRNA had been previously validated for potent knockdown efficiency (51). Following 2 days of doxycycline treatment, HC-04 2B3 cells were analysed by immunoblot for BIM protein expression. The quantity of BIM expression in HC-04 2B3 cells transduced with sgRNA targeting the *BIM* gene was reduced by approximately 75% compared to mock transduced cells, despite not cloning out the population of cells, demonstrating efficient Cas9-mediated loss of BIM expression in these cells (**Figure 1c**). The expression of endogenous BIM remaining was due to either incomplete transduction of HC-04 2B3 cells in the population or to genome repair that was not deleterious to protein expression following Cas9 endonuclease activity, or both. Together, these results demonstrate that HC-04 2B3 were successfully transduced and expressed functional Cas9-mCherry, able to effectively mediate knockdown of BIM protein expression upon transduction with *BIM*-targeting sgRNA. This confirmed the suitability of HC-04 2B3 for broader genome editing applications to study interactions with *P. falciparum* sporozoites.

### Improving upon an established *P. falciparum* cell traversal assay

To enable robust and high-throughput assessment of cell traversal by *P. falciparum* sporozoites, we assessed several parameters with HC-04 2B3 cells. In all cell traversal conditions tested, *P. falciparum* sporozoites dissected from mosquito salivary glands were incubated with hepatocytes for 4 hours to allow traversal to occur. This incubation time was based on our previous finding that cell traversal rates, measured as the percentage of dextran-positive HC-04 cells, increase steadily during the first 2.5 hours post-infection (hpi) but plateaued thereafter (12). First, we evaluated the influence of seeding density on cell traversal. HC-04 2B3 were seeded in 96-well plates at densities ranging from 50,000 cells (∼80%, below confluence) to 100,000 cells (confluent monolayer) and higher densities that formed three-dimensional structures. Although differences were not statistically significant, seeding a confluent monolayer consistently increased cell traversal rates across three independent experiments and two multiplicities of infection (MOIs) (**Figure 2a**, *p*=0.098).

**Figure 2:**
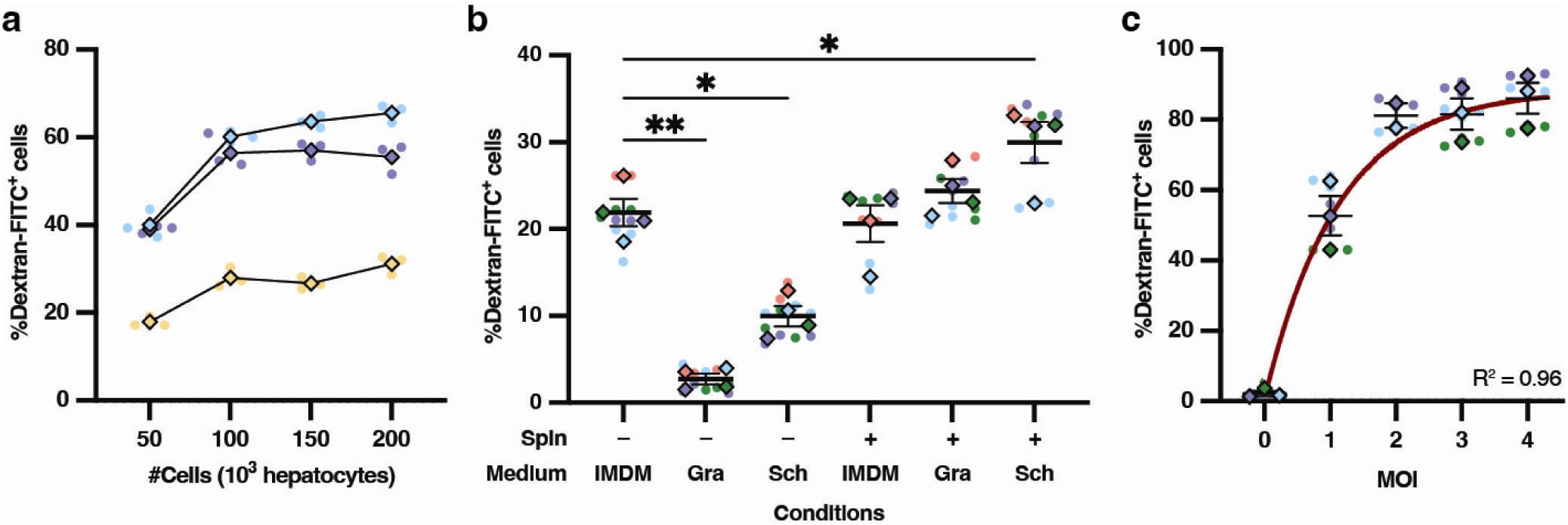
Cell traversal assay optimisation with HC-04 2B3. (**a**) Seeding 100,000 cells led to improved cell traversal rates (however, not significant compared to 50,000 cells; Holm-Sídák multiple comparisons test) (**b**) Dissecting sporozoites into Schneider’s medium followed by centrifugation and resuspension in IMDM significantly improved cell traversal rates (Holm-Sídák multiple comparisons test). When no spin (-) was used, the same medium was used throughout; with spin (+), sporozoites were resuspended in IMDM to support hepatocyte health. (**c**) Increasing the MOI enhanced cell traversal rates, reaching a plateau at 88.4% at MOI 4 (brown curve; exponential plateau fit, R² = 0.96). Superplots (54) present each biological replicate (colour), including technical replicates (round points) and their average (diamond). In panel a, MOI was set at 0.5 instead of 1 for the yellow-coloured replicate. Mean ± SEM are shown in panels b and c. *p < 0.05, **p < 0.01, ***p < 0.001, ****p < 0.0001. Gra, Grace’s Insect medium; Sch, Schneider’s Insect medium.

Next, we assessed how different media and centrifugation of sporozoites impacted viability and cell traversal activity. Dissection and incubation of sporozoites was performed using either Grace’s or Schneider’s insect medium or Iscove’s Modified Dulbecco’s Medium (IMDM). Insect media has been reported to improve sporozoite viability and extend their time of motility (52,53). Dissecting sporozoites into Schneider’s medium, followed by centrifugation and resuspension in IMDM, yielded ideal cell traversal frequencies while also showing that centrifugation did not negatively affect sporozoite (**Figure 2b**, *p* = 0.036 and *p* = 0.56, respectively). Finally, the MOI was increased from the standard 1:1 ratio (one sporozoite per hepatocyte (13)) to 4:1. This resulted in a higher proportion of dextran+ (i.e., traversed) cells. The increase was, however, non-linear and followed an exponential plateau, with a maximum cell traversal efficiency of 92.5% obtained at MOI 4 (**Figure 2c**, *R²*=0.96).

### A refined and harmonized *P. falciparum* cell traversal and invasion assay for screening

With optimized conditions for *P. falciparum* cell traversal established, we next assessed key parameters for measuring productive invasion by sporozoites. Given prior findings and limited consensus in the literature (**Table 1**), we revisited three factors: sporozoite medium, MOI, and the incubation time for sporozoites with hepatocytes in order to evaluate the suitability of these variables for invasion assays, as detailed below.

**Table 1:**
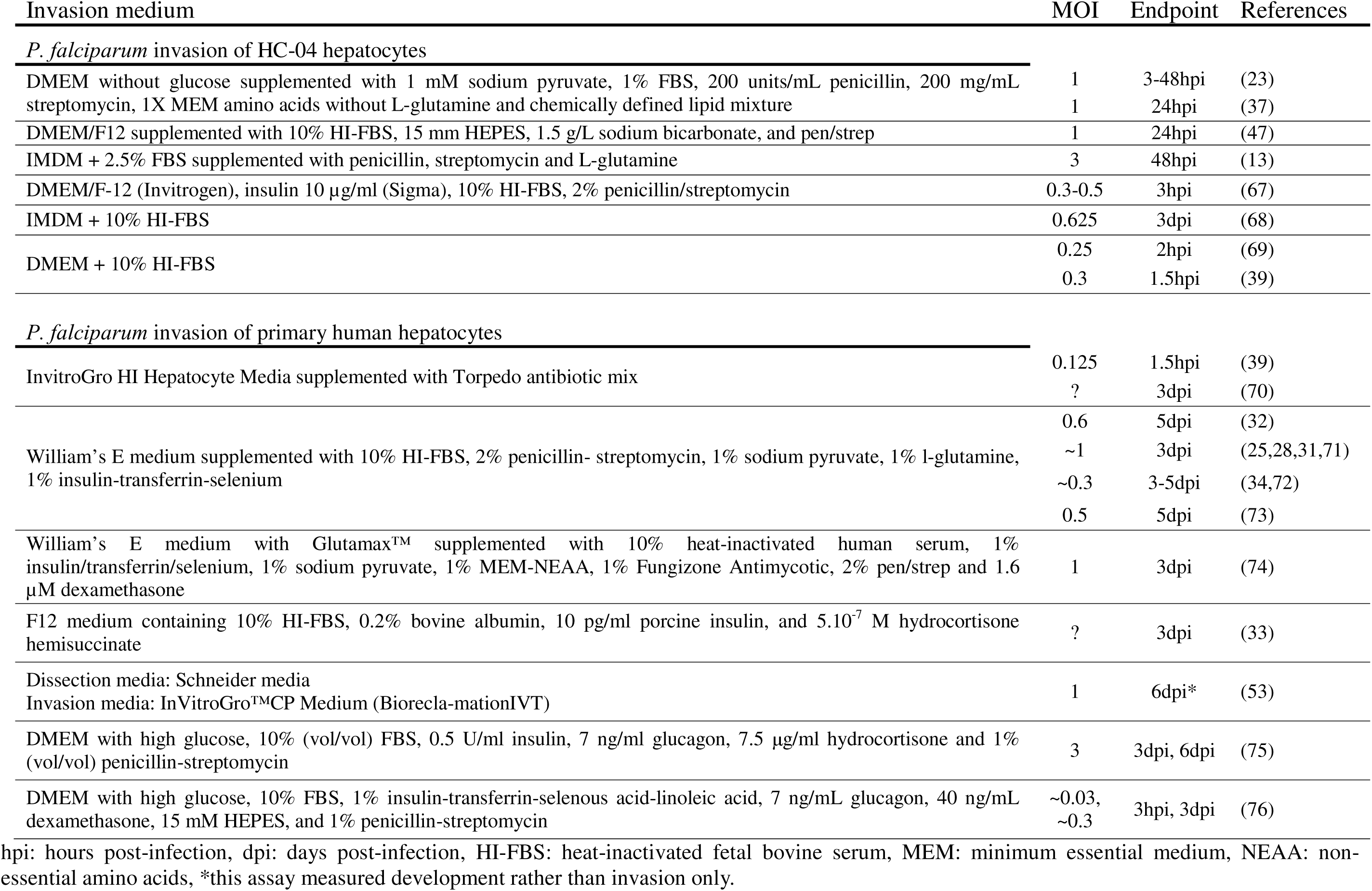
Different parameters for *P. falciparum* sporozoite invasion assays.

Previous studies reported the improved invasion efficiency of HC-04 hepatocytes when *P. falciparum* sporozoites were dissected into the culture medium for mammalian cells, Medium 199 (M199), and then placed into ‘Advanced’ medium, comprising Dulbecco’s Modified Eagle’s Medium (DMEM) with supplements but lacking glucose (see methods), during the invasion assay as this has been reportedly advantageous for *P. falciparum* sporozoite invasion (23,37). Building on our findings regarding optimized cell traversal using insect medium for dissection, we then compared how invasion assay media (culture medium IMDM or ‘Advance’) affected invasion rates after sporozoites were dissected into Schneider’s insect medium. As in the cell traversal assays, we quantified the number of HC-04 2B3 cells containing *P. falciparum* circumsporozoite protein-positive (*Pf*CSP+) parasites after 4 hours, capturing both productive invasion events and non-functional entry (23). There was no significant difference in the percentage of HC-04 2B3 cells containing *Pf*CSP+ parasites between media conditions (**Figure 3a**; *p*>0.05), indicating similar invasion rates when sporozoites were transferred from Schnieder’s into in either IMDM or ‘Advanced’ media. Therefore, subsequent assays involved sporozoite dissection into Schneider’s medium to preserve sporozoite viability, followed by placement into IMDM with serum and co-incubation with hepatocytes to allow sporozoite invasion.

**Figure 3:**
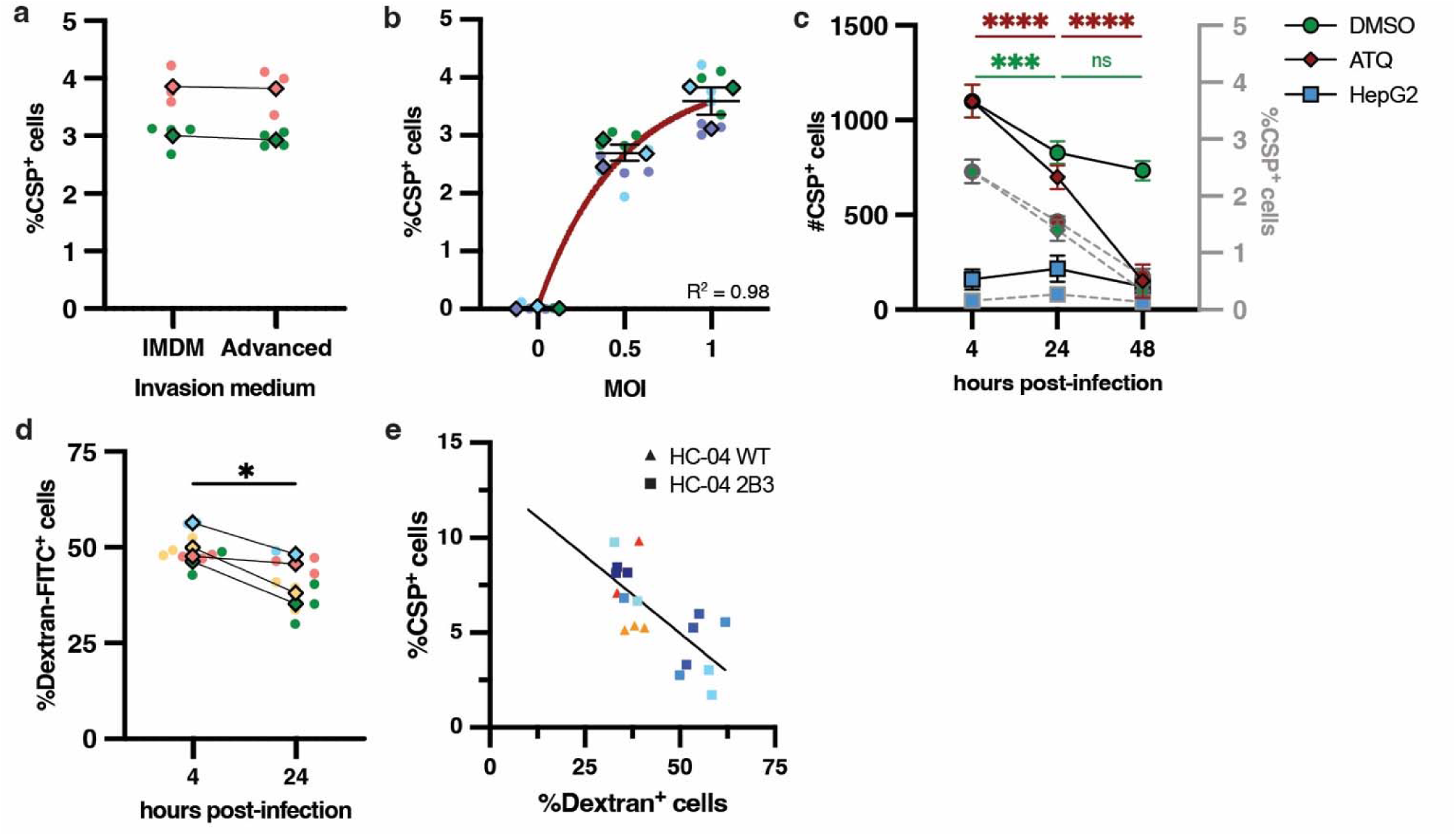
Harmonization of cell traversal and invasion assay parameters for *P. falciparum* sporozoites with HC-04 cells. (**a**) *Pf*CSP+ HC-04 2B3 cells quantification 4 hpi with *P. falciparum* sporozoite isolated in Schneider’s media prior to invasion assay in either IMDM or ‘Advanced’ medium. (*p*=0.2, paired *t*-test). (**b**) Percentage of *Pf*CSP+ HC-04 cells increases with MOI in a non-linear fashion (paired *t*-test, exponential plateau, *R^2^*=0.98). (**c**) Number (black full line and contour) and percentage (grey dashed lines and contour) of *Pf*CSP+ HC-04 2B3 cells across different conditions: ATQ and DMSO are HC-04 cells treated with 50 nM atovaquone or 0.1% DMSO, respectively from 4 hpi. HepG2 were used as a negative control as they are non-permissive to productive invasion by *P. falciparum* sporozoites. Data were comparted using Tukey multiple comparisons test summarised in **Table 1**. (**d**) The percentage of traversed cells significantly decreases between 4h and 24 hpi (paired t-test) from 4 to 24%. (**e**) Matched cell traversal and invasion rates using HC-04 WT (triangle) or HC04 2B3 (square). Cell traversal was quantified 4 hpi, invasion was quantified 24 hpi from the same wells. Colour corresponds to biological replicates, line illustrates significant correlation (Pearson coefficient -0.978, *p*-value=0.004). Superplots (54) in panels **a**-**d** present each biological replicate (colour), including technical replicates (round points) and their average (diamond).

Next, we assessed the effect of MOI on the rate of HC-04 invasion. Since cells that have been traversed are less likely to support infection (55), we focused on an MOI of 1 or lower, as higher MOIs caused higher rates of traversal (**Figure 2c**). As might be expected, incubation of *P. falciparum* sporozoites with HC-04 2B3 cells at MOI 1 resulted in significantly more *Pf*CSP+ host cells than MOI 0.5 (**Figure 3b**, *p*=0.024). Interestingly, while MOI 0.5 consistently resulted in 2-3% *Pf*CSP+ HC-04 cells, doubling the number of sporozoites (MOI 1) did not result in twice the invasion rate measured at 4 hours (**Figure 3b**, exponential plateau, *R^2^*=0.98), suggesting that traversal of the majority of cells impacts negatively invasion (**Figure 2c**, 58% traversal at MOI 1). As such, an MOI 0.5 was selected for subsequent experiments.

Sporozoites either traverse host cells by generating and breaching a TV or invade the cell by forming a PVM within which they develop into exoerythrocytic forms (EEFs). Non-invading parasites are usually cleared within 12 hours (17). With this in mind, we infected HC-04 cells with *P. falciparum* sporozoites and tracked the number of intracellular *Pf*CSP+ parasites for the first 48 hpi. As controls, HepG2 cells that are non-permissive to *P. falciparum* EEF development were included to determine non-functional entry events, as was the antimalarial liver-stage drug atovaquone (ATQ) to kill intracellular EEFs. In untreated controls, there was a reduction in both the number and percentage of *Pf*CSP+ cells between 4 and 24 hpi, consistent with clearance of some EEFs over time, as reported previously (23) which may be due to innate killing of parasites (18,56) (**Figure 3c**, *p*≤0.0005). Interestingly, between 24 and 48 hours, *Pf*CSP+ counts were not significantly different (**Figure 3c**, black contour, *p*=0.32); however, the proportion of *Pf*CSP+ cells decreased markedly due to host cell proliferation that diluted the proportion of infected cells (**Figure 3c**, grey contour *p*<0.0001). On the other hand, ATQ-treatment caused a rapid clearance of infected *Pf*CSP+ HC-04 cells between 4h to 48h as expected (57), with the number of EEFs reaching the same background levels as those in non-permissive HepG2 cells (**Figure 2c**, Tukey multiple comparisons test *p*=0.88). These results indicate that the 24 hpi timepoint was appropriate for distinguishing productive invasion from cell traversal events while mitigating the dilution effect of HC-04 replication evident by 48 hpi.

To study *P. falciparum* sporozoite interactions with hepatocytes, we reasoned it would be advantageous to develop a single assay allowing measurement of cell traversal and productive invasion rates from the same samples. To this end, we measured the cell traversal rates at 4 hpi by quantifying dextran+ cells in half of the experimental wells, while washing the remaining wells to remove extracellular sporozoites and incubating them under normal conditions overnight to allow intracellular parasites to continue their development before quantifying the number of *Pf*CSP+ and dextran+ cells again at 24 hpi. Assuming no significant difference in the number of dextran+ cells across time, this would allow the concomitant measurement of dextran+ (cell traversal) and *Pf*CSP+ (invaded) HC-04 cells at 24 hpi in the future. However, we observed a significant reduction in the number of dextran+ HC-04 cells at 24 hpi when compared to quantification made at 4 hpi, ranging from 4 to 24% (**Figure 3d**, *p*=0.034). This may be explained by HC-04 cell proliferation overnight diluting the dextran signal. We conclude that cell traversal is more accurately quantified on day 1 of the assay (4 hpi) while invasion rates are ideally quantified the following day (24 hpi). We then developed a harmonized assay to simultaneously measure cell traversal and productive invasion from the same samples at 4 and 24 hpi (**Figure 3e**). This integrated approach is valuable given the inverse relationship often observed between traversal and invasion rates and highlights the importance of assessing both processes in parallel to better understand sporozoite-host cell interactions.

### Genetic disruption of 10 human genes involved in infection by diverse pathogens

Having established HC-04 2B3 for the study of *P. falciparum* pre-erythrocytic infection, these cells were genetically modified to generate independent clones deficient in proteins involved in infection by diverse pathogens. We did not test the known *Plasmodium* invasion proteins CD81, EphA2 or SR-B1 due to their tropism for different species (32,35). HC-04 cells also do not express or require CD81 for permissiveness to *P. falciparum* sporozoites (13,28,31), and while the role of EphA2 remains debated (39,58,59), we sought to identify novel factors. Thus, a subset of human genes encoding proteins with biological processes previously reported to be involved in pathogen entry into host cells and/or infection were selected for further study (**Table 2**). Genes were identified based on expression in the liver, location of their encoded proteins on the plasma membrane (or secreted beyond it), and glycosylation enzymes were also included in light of their importance for the conformation, function and surface trafficking of various substrates exploited by pathogens and the role of host proteoglycans in environmental liver sensing by *Plasmodium* sporozoites (37,60–62). Initially, 17 human genes were chosen based on their known or suspected roles in pathogen infection, membrane dynamics, adhesion, endocytosis, and cytoskeletal regulation (**Table 2**). Of these, sgRNAs for 15 genes were present in a Sanger library available to our laboratory (63). Each sgRNA plasmid was picked and amplified for lentiviral particles production. All 15 genes were transduced into HC-04 2B3 in an arrayed format before cloning out with FACS of BFP fluorescence for genotyping and functional analyses of individual clones. Knockout (KO) clones in which the gene of interest was disrupted could not be generated for 5 genes (*RNF223, ESAM, NDST3, NDST1* and *CHIC2*), as the clones did not adhere normally or grew poorly in culture (**Table 2**), and these were not studied further. Clonal KO lines for the remaining 10 genes were successfully generated and validated by next-generation sequencing (**Figure 4a, Supplementary Figure 1**). KO clones were considered validated if they carried a frameshift and/or premature stop codon that prevented expression of the wildtype protein sequence, with preference given to bi-allelic mutations. Altogether, these results showed that 10 independent KO cell line clones were generated by CRISPR/Cas9 genome editing in HC-04 2B3.

**Figure 4:**
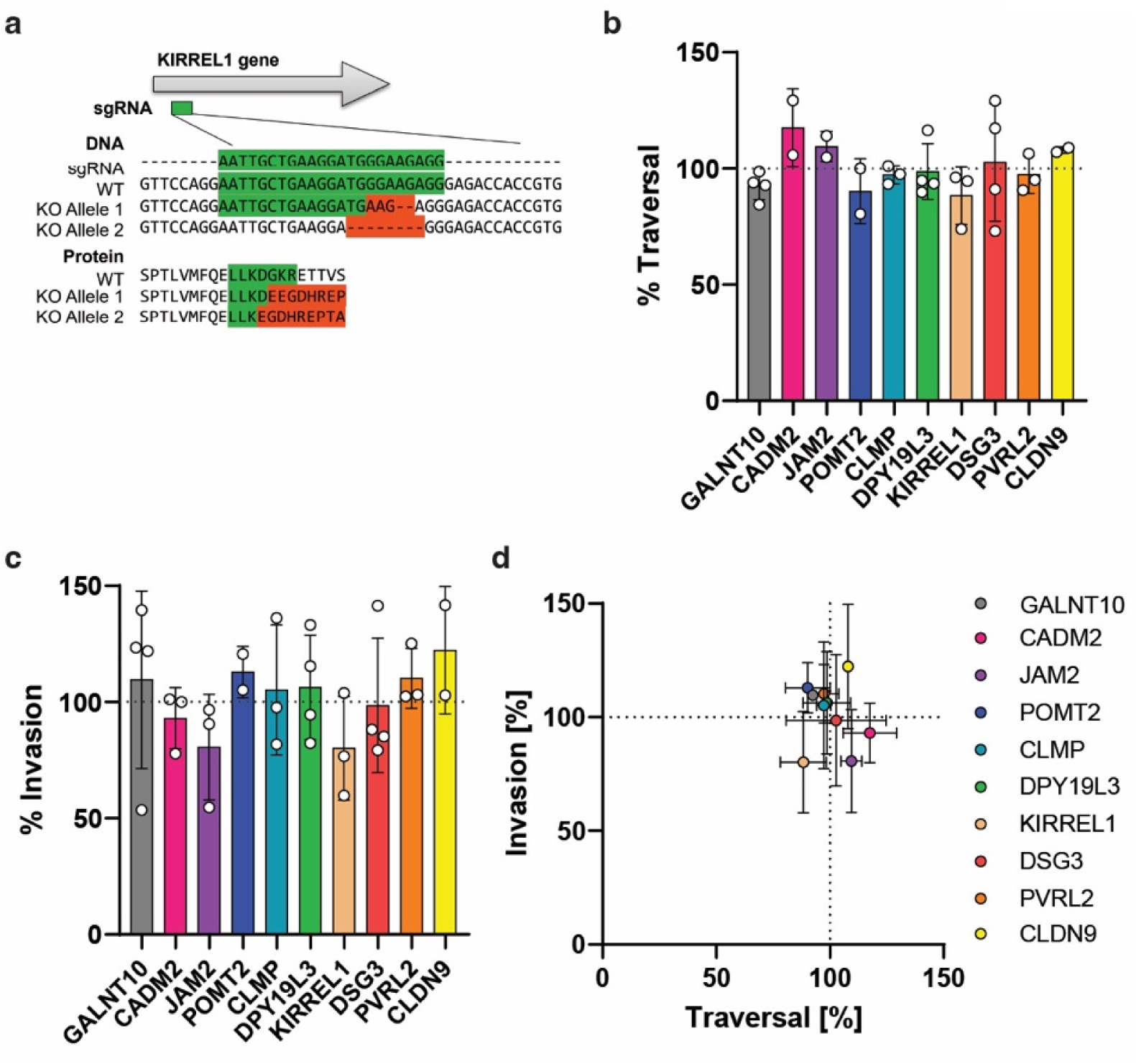
Assessment of *P. falciparum* sporozoite cell traversal and invasion of HC-04 2B3 cell line clones deficient for each of 10 human genes. (**a**) Representative example of a Cas9/sgRNA-mediated genetic disruption of the human *KIRREL1* gene in HC-04 2B3, causing different frameshift and premature STOP codons in each allele. Next generation sequencing of all genes genetically disrupted in individual knockout (KO) clones of 2B3 can be found in Supplementary Figure 1. (**b**) Normalised rates of cell traversal or invasion (**c**) relative to 2B3 parental control cells. No significant reduction was found for any KO cell lines (Kruskal-Wallis test with Dunn’s correction). (**d**) Rates of cell traversal (x-axis) versus invasion (y-axis) from a minimum of 2 biological replicates per human gene KO. Dat are mean +/- SEM. Dashed line represents normalisation to the HC-04 2B3 parental control.

**Table 2:** Selection criteria for forward screening of human genes involved in infection.

|  | Expressed in liver | Present in Sanger library | Successful KO | Microbial infection | Plasma membrane | Extracellular domain | Cell-cell junction | Endo/exocytosis | Lipi metabolism | Immune/inflammation regulation | Significant in previous screen | Secreted | Glycosylation | Pathogen utilising the gene during infection |
| --- | --- | --- | --- | --- | --- | --- | --- | --- | --- | --- | --- | --- | --- | --- |
| *B3GALT5* | N/A | ✗ | N/A | ✗ | ✗ | ✗ | ✗ | ✗ | ✓ | ✓ | ✓ | ✗ | ✓ |  |
| *A4GALT* | ✓ | ✗ | N/A | ✓ | ✗ | ✗ | ✗ | ✗ | ✓ | ✗ | ✓ | ✗ | ✓ | *Bacillus cereus* |
| *POMT2* | ✓ | ✓ | ✓ | ✓ | ✗ | ✗ | ✗ | ✗ | ✗ | ✗ | ✗ | ✗ | ✓ | Different viruses |
| *GALNT10* | ✓ | ✓ | ✓ | ✓ | ✗ | ✗ | ✗ | ✗ | ✗ | ✗ | ✓ | ✗ | ✓ | *Salmonella* |
| *RNF223* | N/A | ✓ | ✗ | ✗ | ✓ | N/A | ✗ | ✗ | ✗ | ✗ | ✓ | ✗ | ✗ |  |
| *ESAM* | ✓ | ✓ | ✗ | ✗ | ✓ | ✓ | ✓ | ✗ | ✗ | ✓ | ✓ | ✗ | ✗ |  |
| *NDST3* | ✓ | ✓ | ✗ | ✗ | ✗ | ✗ | ✗ | ✗ | ✗ | ✓ | ✓ | ✗ | ✗ |  |
| *NDST1* | ✓ | ✓ | ✗ | ✓ | ✗ | ✗ | ✗ | ✓ | ✗ | ✓ | ✓ | ✗ | ✗ | Poxvirus, HS modulation |
| *CHIC2* | ✓ | ✓ | ✗ | ✗ | ✓ | ✗ | ✗ | ✓ | ✗ | ✓ | ✗ | ✗ | ✗ | HS modulation |
| *CELA2A* | ✓ | ✓ | ✓ | ✗ | ✗ | ✗ | ✗ | ✗ | ✗ | ✓ | ✓ | ✓ | ✗ |  |
| *KIRREL1* | ✓ | ✓ | ✓ | ✗ | ✓ | ✓ | ✓ | ✗ | ✗ | ✗ | ✓ | ✗ | ✗ |  |
| *CLMP* | ✓ | ✓ | ✓ | ✗ | ✓ | ✓ | ✓ | ✗ | ✗ | ✗ | ✗ | ✗ | ✗ |  |
| *DSG3* | N/A | ✓ | ✓ | ✗ | ✓ | ✓ | ✓ | ✗ | ✗ | ✗ | ✗ | ✗ | ✗ |  |
| *PVRL2* | ✓ | ✓ | ✓ | ✓ | ✓ | ✓ | ✓ | ✓ | ✗ | ✓ | ✗ | ✗ | ✗ | Poliovirus |
| *CDLN9* | ✓ | ✓ | ✓ | ✓ | ✓ | ✓ | ✓ | ✓ | ✗ | ✗ | ✗ | ✗ | ✗ | Hepatitis C virus |
| *JAM2* | ✓ | ✓ | ✓ | ✓ | ✓ | ✓ | ✓ | ✓ | ✗ | ✓ | ✗ | ✗ | ✗ | Orthoreoviruses |
| *CADM2* | ✓ | ✓ | ✓ | ✓ | ✓ | ✓ | ✓ | ✗ | ✗ | ✗ | ✓ | ✗ | ✗ | Measles virus |

Host candidate genes and their association with functional categories. Each column indicates whether a gene is involved in the listed pathway or feature. **Green tick**, direct involvement; **yellow tick**, indirect involvement; **red cross**, no known relation. “Significant in previous screen” refers to hits from the *P. yoelii* CRISPR screen (58). “HS” = heparan sulfates. The final column lists pathogens reported to utilize the indicated gene product during infection.

### Characterization of *P. falciparum* sporozoite infectivity for HC-04 2B3 KO cell line clones

To assess whether any of the 10 gene candidates are important for *P. falciparum* sporozoite-infectivity, the rates of cell traversal and productive invasion was measured in the harmonized cytometric assay developed above. Initially, 10 KO cell line clones were selected for assessment and results were compared to the parental HC-04 2B3 clone in an arrayed format (**Figure 4b-d**). Though none of the KO clones showed a significant reduction in cell traversal or invasion by *P. falciparum* sporozoites, all cell lines demonstrated an inverse correlation between the rate of cell traversal and the rate of invasion (**Figure 4d**, upper left or lower right panel), though *KIRREL1* showed a slight decrease in both cell traversal and invasion. The absence of a significant effect on cell traversal or invasion suggests that the 10 genes selected are not crucial for these processes, although we cannot exclude the possibility that genes may have functional redundancy with other unknown host factors that could mask a defect.

## Discussion

The molecular interplay between *P. falciparum* sporozoites and their host hepatocytes are important for the outcome of infection yet incompletely understood. In this study, we engineered the tools necessary to begin dissecting the host determinants of infection by the human malaria parasite at scale. By generating and validating HC-04 2B3 with functional Cas9-mCherry endonuclease activity, we enabled precise genetic manipulation of the only human cell line known to support *P. falciparum* liver-stage development. HC-04 2B3 demonstrated efficient Cas9 activity and retained normal susceptibility to sporozoite traversal and invasion, confirming the utility of these hepatocytes for functional studies with human malaria parasites.

We refined key assay parameters to improve reproducibility and sensitivity. Notably, decoupling mosquito dissection and assay media proved valuable: sporozoite traversal efficiency increased significantly when mosquitoes were dissected in insect medium rather than culture medium, consistent with prior reports on extended sporozoite viability in *P. vivax* (52,53). This suggests that the benefit of ‘Advanced’ medium described previously may reflect improved sporozoite preservation rather than true enhancement of hepatocyte infection (37). Additionally, the rates of cell traversal were more consistent when using confluent HC-04 monolayers, suggesting this may better mimic the physiological microenvironment. We also evaluated how the MOI input influences the rate of cell traversal and invasion, with a plateau occurring at a *P. falciparum*-to-HC-04 ratio above 3. A small but consistent subset of cells (>5%) remained dextran-negative, raising the possibility that resistance to traversal by *P. falciparum* may occur by some cells of the population.

The rate of HC-04 invasion by sporozoites plateaued at lower MOIs compared to cell traversal, perhaps due to cell traversal-induced changes to the cell that reduced cellular susceptibility, such as NF-κB activation that was reported previously (55). These findings support the need in future genetic screens to find a balance between a maximal MOI for cell traversal that preserves biological relevance for invasion while maximising assay sensitivity given the limited availability of sporozoites that are dissected from individual mosquitoes in each experiment. Temporal resolution also proved important for measuring parasite infectivity. While intracellular staining at 4 hpi captures both cell traversal and invaded parasites, this time point cannot clearly distinguish between the two events. By 24 hpi however, non-viable sporozoites are typically cleared (16,17), providing a clearer snapshot of productive invasion that occurred the day before, while limiting confounding effects from cell proliferation on measuring cell traversal on the second day. Although sporozoites can abort traversal and lodge inside the nucleus and persist for some time, as described with HepG2 cells (28), these events are rare and we believe unlikely to affect the results measured in the harmonized cytometric assay we developed. As such, we conclude that 24 hpi represents a reliable endpoint for quantifying *P. falciparum* sporozoite invasion but is unsuitable for assessing the rate of cell traversal.

Our harmonised assay, which allows concurrent measurement of cell traversal and invasion in the same sample at distinct timepoints, increases efficiency and reduces sporozoite usage by half. This integrated approach is particularly useful given the often inverse relationship observed between traversal and invasion (55), where high cell traversal can reduce productive invasion (**Figure 3e**). As such, simultaneous assessment of both processes is essential for accurately capturing the complexity of host–parasite interactions.

Applying these conditions, we performed the first arrayed CRISPR-Cas9 screen of a small number of human genes exploited by diverse pathogens for a role in *P. falciparum* sporozoite infectivity. None of the genes that we disrupted successfully in this study were critical for *P. falciparum* cell traversal or invasion using the assays we developed, though they may be involved in downstream pathways essential in EEF development that our short-term assay does not allow us to interrogate. Additionally, specific characteristics of HC-04 cells might impact the results, as differences in productive invasion between primary human hepatocytes and HC-04 cells have been reported (13,28,29). Yet this work provides proof-of-concept of the utility of HC-04 2B3 for conducting whole genome CRISPR screens to dissect different *P. falciparum* phenotypes at the liver stage.

A major limitation for whole genome screening is the requirement for high numbers of sporozoites that must be dissected from the salivary glands of hundreds of individual mosquitoes just prior to the experiment. This limits the MOI for the screen, which may constrain the feasibility or study design of genome-wide liver-stage screens for malaria. However, targeted libraries enriched for hepatocyte-specific, surface-expressed, or infection-responsive genes represent a strategic and increasingly tractable alternative to assessing focussed genome subsets in individual screens. With continued optimization, including improved sporozoite production, miniaturisation of assay formats, and automation of readouts, this platform provides a strong foundation for high-throughput discovery of host-determinants of *P. falciparum* hepatocyte infection. Beyond genome screens, HC-04 2B3 are well suited for mechanistic studies of early host–parasite interactions such as membrane repair, innate immune evasion, and receptor engagement and intracellular development. Ultimately, the *in vitro* assays developed in this study can enable systematic dissection of the host landscape exploited by *P. falciparum* during liver-stage infection.

Together, our study underscores the value of engineered HC-04 2B3 cell lines and refined phenotypic assays as foundational tools for human liver-stage malaria research. Future efforts should aim to extend this platform to longer-term assays for exoerythrocytic form (EEF) development, combinatorial gene knockouts, or larger pooled CRISPR screens to uncover the complex network of host factors influencing *P. falciparum* liver-stage biology. HC-04 2B3 cells are also a valuable tool for the study of hepatocyte biology and infection dynamics by liver-tropic pathogens.

## Material and Methods

### Cell maintenance and lentivirus production

HC-04 hepatocytes (46) were routinely cultured in Iscove’s Modified Dulbecco’s Medium (IMDM, Life Technologies Cat#12200) supplemented with 5% of foetal bovine serum (FBS) and penicillin and streptomycin (pen/strep). HEK293T cells were maintained in Dulbecco’s Modified Eagle Medium (DMEM, Life Technologies Cat#31600) complemented with 10% FBS and pen/strep. Both cell lines were passaged every 2-4 days and cultured for no more than 2 months.

### Lentiviral production, transduction and virus titration

Lentivirus production was performed as previously described (64). Briefly, HEK293T were seeded 24h prior to calcium phosphate co-transfection of target DNA with envelope, packaging, replication plasmids (from Didier Trono, Addgene#12251, Addgene#12253 and Addgene#12259 respectively, (65)). For Cas9, FUCas9Cherry plasmid was used as target DNA (Addgene # 70182, (51)). For the individual genes knock-outs, gene-specific sgRNAs (N=2) derived from the Sanger library were used as target DNA (63). Cells were incubated overnight, and medium was changed the following morning. Both 24h and 48h supernatants were collected, filtered through 0.45µm and stored at -80°C until further use.

HC-04 cells were seeded at 100,000 cells per well in 6-well plates the day prior to transduction. For Cas9 transduction, 5 mL of lentiviral supernatant containing 2 µg/mL of polybrene were added and particles were centrifuged onto cells at 2,200rpm for 2h at 35°C. Cells were washed off twice and incubated 24h before a second transduction was performed. Similarly, sgRNA transduction of individual genes were performed by a single transduction using 2 mL of lentiviral supernatant containing 2 µg/mL of polybrene. Cells were washed twice daily for the first 2 days. In all cases, transduction was confirmed two days later by confirming fluorescence on BD LSRFortessa™ X-20 flow cytometer. Clonal cell lines were established by sorting individual fluorescent cells using a BD FACSAria flow cytometer, Cas9 clones being mCherry-fluorescent for and target gene KO cell lines BFP-positive.

### Cas9 activity by immunoblotting

Cas9 activity was confirmed by immunoblotting. HC-04 Cas9+ 2B3 clone were transduced with the *isgBIM* huEx3 construct (51) targeting BIM peptide with a doxycycline-inducible promoter or an empty plasmid. Two days post-transduction, transduced cells were split into two populations, one which was incubated with 1 µg/mL of doxycycline while the control stayed in culture medium for another two days. Cells were then trypsinized, washed twice with DPBS and dry pellets were frozen at -80°C. Immunoblotting was performed following (51) with minor adjustments. Briefly, cells were lysed in RIPA buffer containing 1X protease and phosphatase inhibitors for 30min on ice. DNA material was pelleted at 10,000 x g for 5 min at 4°C. Total protein extracts were denatured at 96°C for 5 min and were separated on SDS-PAGE gel 4-12% polyacrylamide (Invitrogen NP0321). After transfer onto a 0.45 μm nitrocellulose membrane (Amersham), samples were blocked and probed (anti-BIM, Enzo Life Sciences ADI-AAP-330-E and anti-β-actin clone AC74, Sigma-Aldrich kindly provided by Dr Gemma Kelly, WEHI; anti-rabbit and anti-mouse IgG-HRP, Invitrogen) in 5% milk/Dulbecco’s phosphate buffered saline. Imaging was performed with either ECL Western Blotting Detection Reagent kit or the ECL Prime kit (Amersham) depending on the signal. Super RX x-ray films (Fuji Film) were developed by Kodak X-OMAT processor.

### *P. falciparum* maintenance and gametocyte induction

*P. falciparum* NF54 strain was cultured in O-positive human blood (obtained from Melbourne Red Cross) as described before (12). Early parasites were synchronized using regular sorbitol treatment. Synchronous culture was used for gametocyte induction which was performed following the crash method (66), with daily medium change for 16 consecutive days.

### Mosquito rearing and infection

*Anopheles stephensi* mosquitoes were infected with *P. falciparum* gametocytes (0.3% stage V gametocytaemia, 50% hematocrit) 1-3 days post-enclosure. They were deprived of sugar for 48h to ensure only the fittest engorged females were conserved for subsequent dissection. Oocyst counts were performed 7 days post-infection using mercurochrome staining. Sporozoite dissections took place 14-18 days post-infection.

### Cell traversal assay

HC-04 cells were passaged, and 100,000 cells (or specified number) were seeded per well of a 96-well plate the day before the assay. Salivary gland dissections were carried out in Schneider’s insect medium (unless otherwise stated) and lasted no more than 1h. Sporozoites were released and filtered through glass wool before count. They were pelleted for 3 min at 10,000 x g at 4°C and resuspended in IMDM containing 10% heat-inactivated human serum (HIHS), 0.5 mg/mL dextran-FITC (Invitrogen D1821). After addition to cells, parasites were pelleted onto hepatocytes at 100 x g for 3 min and incubated for 4h at 37°C. Cell traversal rates were analysed by flow cytometry (BD LSRFortessa™ X-20 flow cytometer) after cells were trypsinized, washed and resuspended in DPBS.

### Invasion assay

One day prior the assay, 100,000 HC-04 cells were seeded per well in a 96-well plate. Dissection and sporozoite isolation were performed as described above, and they were resuspended in IMDM or ‘Advanced’ medium (DMEM without glucose (Life Technologies, 11966-025) containing 1% active FBS (Cellgro, 35-010-CV), 1 mM sodium pyruvate (Life Technologies, 11360-070), 1X MEM non-essential amino acids without L-glutamine (Sigma-Aldrich, M5550), 1:500 dilution of Lipid Mixture 1, Chemically Defined (Sigma-Aldrich, L0-288) and 1X Pen/Strep (Corning, 30-001-Cl) (37)) supplemented with 10% HIHS for an MOI of 0.5. Plates were centrifuged at 100 x g for 3 min and incubated for 4h. Cells were washed once in DPBS and, for assays beyond 4h, cultured in IMDM supplemented with 5% FBS, 2.5 µg/mL amphotericin B, 110 µM neomycin and 50 µg/mL gentamicin. At the designated endpoint, cells were trypsinized, fixed and stained using Cytoperm™ kit (BD Biosciences, 554714) as previously described (39). In short, fixation was performed in Cytofix for 15 min on ice, followed by two washes and staining in 1× Cytoperm for 1 hour with 1 µg/mL of α-PfCSP 2A10 monoclonal antibody (MR4, MRA-183A) conjugated to Alexa Fluor 647. Cells were washed twice in Cytoperm solution before being resuspended in DPBS and intracellular parasites were measured by flow cytometry (BD LSRFortessa™ X-20 flow cytometer).

### Harmonised cell traversal and invasion assay for screening

Harmonised assays were performed as cell traversal with the following adjustments. After 4h incubation at 37°C, cells were washed with DPBS and trypsinized for 5 min. Wells were resuspended in 200 µL of IMDM supplemented with 5% FBS, 2.5 µg/mL amphotericin B, 110 µM neomycin and 50 µg/mL gentamicin and 100 µL was transferred in a fresh 96-well round bottom. This new plate was centrifugated at 300 x g for 3 min, and cells were resuspended in DPBS before cell traversal assessment by flow cytometry (BD LSRFortessa™ X-20 flow cytometer). The culture plate was incubated at 37°C for another 20h after which hepatocyte invasion was determined as described above.

### Next Generation Sequencing

Genetic disruption of target genes was confirmed by next-generation sequencing following overhang-based PCR preparation. Clonal cell lines were generated, and gene-specific primers were designed to flank target regions, producing 250–300 bp amplicons (300-cycle kit). Each primer included a universal 5′ sequencing overhang: Forward 5′-GTGACCTATGAACTCAGGAGTC-3′; Reverse 5′-CTGAGACTTGCACATCGCAGC-3′. PCR was performed in triplicate in 96-well format with a no-template control per primer pair. Each 20 µL reaction contained 1 µL genomic DNA (∼100 ng/µL), 10 µL GoTaq Green Mix, 0.5 µL of each primer (10 µM), and 8 µL H₂O. Cycling conditions: 95 °C for 3 min; 18 cycles of 95 °C for 15 s, 60 °C for 30 s, 72 °C for 30 s; final extension at 72 °C for 7 min; hold at 10 °C. PCR products were cleaned using 20 µL (1:1) NGS beads, incubated 5 min, and separated on a magnetic rack. Supernatant was removed, and wells were washed twice with 150 µL 70% ethanol. After air drying, DNA was eluted in 40 µL H₂O, and 30 µL transferred to a new 96-well plate.

For indexing PCR, uniquely barcoded overhang primers (10 µM) were arranged so each clone received a unique primer pair. Each 20 µL reaction included 10 µL cleaned DNA, 10 µL GoTaq Green Mix, and 0.5 µL of each indexing primer. Cycling: 95 °C for 3 min; 24 cycles of 95 °C for 15 s, 60 °C for 30 s, 72 °C for 30 s; final extension at 72 °C for 7 min; hold at 10 °C. Reactions were spot-checked via agarose gel electrophoresis. For pooling, 5 µL from each well (including controls) were combined, mixed in a 25 mL reservoir, and transferred to a 1.5 mL tube. From this pool, 50 µL underwent bead cleanup using 40 µL NGS beads. After 5 min, beads were collected, washed twice with 180 µL ethanol, air-dried, and eluted in 105 µL H₂O. After 5 min, beads were cleared on a magnetic rack, and the eluate was used for sequencing. Libraries were sequenced on an Illumina MiSeq, and data were analyzed in DNASTAR MegAlign Pro by aligning reads to the HC-04 2B3 parent clone and respective sgRNA to confirm gene disruptions.

## Acknowledgements

The authors would like to thank the Melbourne Red Cross for human erythrocytes, Melissa Hobbs and Dr Julie Healer for their support in mosquito rearing and insectary management, Dr Amber Alsop and A/Prof Kym Lowes for their support in accessing the Sanger library, Prof Gemma Kelly for kindly providing us with antibodies, and Dr Stephen Wilcox for providing support for NGS of our clonal cell lines. We would like to extend our thanks to the members of the Boddey laboratory for their support and advice, in particular Dr Ryan Steel and Sash Lopaticki for substantial assistance and insightful discussions on host-parasite interactions. This work was funded by the National Health & Medical Research Council of Australia (GNT1176955) and supported by the Victorian State Government Operational Infrastructure Support grant (Institutional grant) and Australian Government NHMRC IRIISS.

## Author contributions

E.H., L.H.V., and J.A.B. conceptualized the studies and wrote the draft, and all authors contributed to data analysis and reviewing and editing the manuscript. E.H., L.H.V., and M.D. performed biological experiments.

## References

1. World malaria report 2024. Geneva, Switzerland: World Health Organization; p. 316.

2. Loubens M, Vincensini L, Fernandes P, Briquet S, Marinach C, Silvie O. Plasmodium sporozoites on the move: Switching from cell traversal to productive invasion of hepatocytes. Mol Microbiol. 2021 May;115(5):870–81.

3. Arredondo SA, Schepis A, Reynolds L, Kappe SHI. Secretory Organelle Function in the Plasmodium Sporozoite. Trends Parasitol. 2021 July;37(7):651–63.

4. Frischknecht F, Matuschewski K. Plasmodium Sporozoite Biology. Cold Spring Harb Perspect Med. 2017 May 1;7(5):a025478.

5. Douglas RG, Moon RW, Frischknecht F. Cytoskeleton Organization in Formation and Motility of Apicomplexan Parasites. Annu Rev Microbiol. 2024 Nov;78(1):311–35.

6. Yang ASP, Boddey JA. Molecular mechanisms of host cell traversal by malaria sporozoites. Int J Parasitol. 2017 Feb;47(2–3):129–36.

7. Vanderberg JP, Chew S, Stewart MJ. Plasmodium sporozoite interactions with macrophages in vitro: a videomicroscopic analysis. J Protozool. 1990;37(6):528–36.

8. Mota MM, Pradel G, Vanderberg JP, Hafalla JC, Frevert U, Nussenzweig RS, et al. Migration of Plasmodium sporozoites through cells before infection. Science. 2001 Jan 5;291(5501):141–4.

9. Ishino T, Yano K, Chinzei Y, Yuda M. Cell-passage activity is required for the malarial parasite to cross the liver sinusoidal cell layer. PLoS Biol. 2004 Jan;2(1):E4.

10. Amino R, Giovannini D, Thiberge S, Gueirard P, Boisson B, Dubremetz JF, et al. Host cell traversal is important for progression of the malaria parasite through the dermis to the liver. Cell Host Microbe. 2008 Feb 14;3(2):88–96.

11. Tavares J, Formaglio P, Thiberge S, Mordelet E, Van Rooijen N, Medvinsky A, et al. Role of host cell traversal by the malaria sporozoite during liver infection. J Exp Med. 2013 May 6;210(5):905–15.

12. Yang ASP, O’Neill MT, Jennison C, Lopaticki S, Allison CC, Armistead JS, et al. Cell Traversal Activity Is Important for Plasmodium falciparum Liver Infection in Humanized Mice. Cell Rep. 2017 Mar 28;18(13):3105–16.

13. Dumoulin PC, Trop SA, Ma J, Zhang H, Sherman MA, Levitskaya J. Flow Cytometry Based Detection and Isolation of Plasmodium falciparum Liver Stages In Vitro. PloS One. 2015;10(6):e0129623.

14. Thiberge S, Blazquez S, Baldacci P, Renaud O, Shorte S, Ménard R, et al. In vivo imaging of malaria parasites in the murine liver. Nat Protoc. 2007;2(7):1811–8.

15. Frevert U, Engelmann S, Zougbédé S, Stange J, Ng B, Matuschewski K, et al. Intravital observation of Plasmodium berghei sporozoite infection of the liver. PLoS Biol. 2005 June;3(6):e192.

16. Risco-Castillo V, Topçu S, Marinach C, Manzoni G, Bigorgne AE, Briquet S, et al. Malaria Sporozoites Traverse Host Cells within Transient Vacuoles. Cell Host Microbe. 2015 Nov 11;18(5):593–603.

17. Bindschedler A, Wacker R, Egli J, Eickel N, Schmuckli-Maurer J, Franke-Fayard BM, et al. Plasmodium berghei sporozoites in nonreplicative vacuole are eliminated by a PI3P-mediated autophagy-independent pathway. Cell Microbiol. 2021 Jan;23(1):e13271.

18. Marques-da-Silva C, Schmidt-Silva C, Bowers C, Charles-Chess NAE, Samuel C, Shiau JC, et al. Type I interferons induce guanylate-binding proteins and lysosomal defense in hepatocytes to control malaria. Cell Host Microbe. 2025 Apr 9;33(4):529–544.e9.

19. Ishino T, Chinzei Y, Yuda M. A Plasmodium sporozoite protein with a membrane attack complex domain is required for breaching the liver sinusoidal cell layer prior to hepatocyte infection. Cell Microbiol. 2005 Feb;7(2):199–208.

20. Kariu T, Ishino T, Yano K, Chinzei Y, Yuda M. CelTOS, a novel malarial protein that mediates transmission to mosquito and vertebrate hosts. Mol Microbiol. 2006 Mar;59(5):1369–79.

21. Steel RWJ, Pei Y, Camargo N, Kaushansky A, Dankwa DA, Martinson T, et al. Plasmodium yoelii S4/CelTOS is important for sporozoite gliding motility and cell traversal. Cell Microbiol. 2018 Apr;20(4).

22. Jimah JR, Salinas ND, Sala-Rabanal M, Jones NG, Sibley LD, Nichols CG, et al. Malaria parasite CelTOS targets the inner leaflet of cell membranes for pore-dependent disruption. eLife. 2016 Dec 1;5:e20621.

23. Yang ASP, Lopaticki S, O’Neill MT, Erickson SM, Douglas DN, Kneteman NM, et al. AMA1 and MAEBL are important for Plasmodium falciparum sporozoite infection of the liver. Cell Microbiol. 2017 Sept;19(9).

24. Angage D, Chmielewski J, Maddumage JC, Hesping E, Caiazzo S, Lai KH, et al. A broadly cross-reactive i-body to AMA1 potently inhibits blood and liver stages of Plasmodium parasites. Nat Commun. 2024 Aug 22;15(1):7206.

25. Silvie O, Franetich JF, Charrin S, Mueller MS, Siau A, Bodescot M, et al. A role for apical membrane antigen 1 during invasion of hepatocytes by Plasmodium falciparum sporozoites. J Biol Chem. 2004 Mar 5;279(10):9490–6.

26. Gonzalez V, Combe A, David V, Malmquist NA, Delorme V, Leroy C, et al. Host cell entry by apicomplexa parasites requires actin polymerization in the host cell. Cell Host Microbe. 2009 Mar 19;5(3):259–72.

27. Vijayan K, Cestari I, Mast FD, Glennon EKK, McDermott SM, Kain HS, et al. Plasmodium Secretion Induces Hepatocyte Lysosome Exocytosis and Promotes Parasite Entry. iScience. 2019 Nov 22;21:603–11.

28. Silvie O, Rubinstein E, Franetich JF, Prenant M, Belnoue E, Rénia L, et al. Hepatocyte CD81 is required for Plasmodium falciparum and Plasmodium yoelii sporozoite infectivity. Nat Med. 2003 Jan;9(1):93–6.

29. Foquet L, Hermsen CC, Verhoye L, van Gemert GJ, Cortese R, Nicosia A, et al. Anti-CD81 but not anti-SR-BI blocks Plasmodium falciparum liver infection in a humanized mouse model. J Antimicrob Chemother. 2015;70(6):1784–7.

30. Risco-Castillo V, Topçu S, Son O, Briquet S, Manzoni G, Silvie O. CD81 is required for rhoptry discharge during host cell invasion by Plasmodium yoelii sporozoites. Cell Microbiol. 2014 Oct;16(10):1533–48.

31. Silvie O, Greco C, Franetich JF, Dubart-Kupperschmitt A, Hannoun L, van Gemert GJ, et al. Expression of human CD81 differently affects host cell susceptibility to malaria sporozoites depending on the Plasmodium species. Cell Microbiol. 2006 July;8(7):1134–46.

32. Manzoni G, Marinach C, Topçu S, Briquet S, Grand M, Tolle M, et al. Plasmodium P36 determines host cell receptor usage during sporozoite invasion. eLife. 2017 May 16;6:e25903.

33. Rodrigues CD, Hannus M, Prudêncio M, Martin C, Gonçalves LA, Portugal S, et al. Host scavenger receptor SR-BI plays a dual role in the establishment of malaria parasite liver infection. Cell Host Microbe. 2008 Sept 11;4(3):271–82.

34. Yalaoui S, Huby T, Franetich JF, Gego A, Rametti A, Moreau M, et al. Scavenger receptor BI boosts hepatocyte permissiveness to Plasmodium infection. Cell Host Microbe. 2008 Sept 11;4(3):283–92.

35. Langlois AC, Manzoni G, Vincensini L, Coppée R, Marinach C, Guérin M, et al. Molecular determinants of SR-B1-dependent Plasmodium sporozoite entry into hepatocytes. Sci Rep. 2020 Aug 11;10(1):13509.

36. Raphemot R, Toro-Moreno M, Lu KY, Posfai D, Derbyshire ER. Discovery of Druggable Host Factors Critical to Plasmodium Liver-Stage Infection. Cell Chem Biol. 2019 Sept 19;26(9):1253–1262.e5.

37. Tweedell RE, Tao D, Hamerly T, Robinson TM, Larsen S, Grønning AGB, et al. The Selection of a Hepatocyte Cell Line Susceptible to Plasmodium falciparum Sporozoite Invasion That Is Associated With Expression of Glypican-3. Front Microbiol. 2019;10:127.

38. Amanzougaghene N, Tajeri S, Yalaoui S, Lorthiois A, Soulard V, Gego A, et al. The Host Protein Aquaporin-9 is Required for Efficient Plasmodium falciparum Sporozoite Entry into Human Hepatocytes. Front Cell Infect Microbiol. 2021;11:704662.

39. Kaushansky A, Douglass AN, Arang N, Vigdorovich V, Dambrauskas N, Kain HS, et al. Malaria parasites target the hepatocyte receptor EphA2 for successful host infection. Science. 2015 Nov 27;350(6264):1089–92.

40. Mittal N, Davis C, McLean P, Calla J, Godinez-Macias KP, Gardner A, et al. Human nuclear hormone receptor activity contributes to malaria parasite liver stage development. Cell Chem Biol. 2023 May 18;30(5):486–498.e7.

41. Hesping E, Boddey JA. Whole-genome CRISPR screens to understand Apicomplexan-host interactions. Mol Microbiol. 2024 Apr;121(4):717–26.

42. Gibson AR, Sateriale A, Dumaine JE, Engiles JB, Pardy RD, Gullicksrud JA, et al. A genetic screen identifies a protective type III interferon response to Cryptosporidium that requires TLR3 dependent recognition. PLoS Pathog. 2022 May;18(5):e1010003.

43. Doerflinger M, Forsyth W, Ebert G, Pellegrini M, Herold MJ. CRISPR/Cas9-The ultimate weapon to battle infectious diseases? Cell Microbiol. 2017 Feb;19(2).

44. Vijayan K, Arang N, Wei L, Morrison R, Geiger R, Parks KR, et al. A genome-wide CRISPR-Cas9 screen identifies CENPJ as a host regulator of altered microtubule organization during Plasmodium liver infection. Cell Chem Biol. 2022 Sept 15;29(9):1419–1433.e5.

45. Hollingdale MR, Nardin EH, Tharavanij S, Schwartz AL, Nussenzweig RS. Inhibition of entry of Plasmodium falciparum and P. vivax sporozoites into cultured cells; an in vitro assay of protective antibodies. J Immunol Baltim Md 1950. 1984 Feb;132(2):909–13.

46. Sattabongkot J, Yimamnuaychoke N, Leelaudomlipi S, Rasameesoraj M, Jenwithisuk R, Coleman RE, et al. Establishment of a human hepatocyte line that supports in vitro development of the exoerythrocytic stages of the malaria parasites Plasmodium falciparum and P. vivax. Am J Trop Med Hyg. 2006 May;74(5):708–15.

47. Tao D, King JG, Tweedell RE, Jost PJ, Boddey JA, Dinglasan RR. The acute transcriptomic and proteomic response of HC-04 hepatoma cells to hepatocyte growth factor and its implications for Plasmodium falciparum sporozoite invasion. Mol Cell Proteomics MCP. 2014 May;13(5):1153–64.

48. VanBuskirk KM, O’Neill MT, De La Vega P, Maier AG, Krzych U, Williams J, et al. Preerythrocytic, live-attenuated Plasmodium falciparum vaccine candidates by design. Proc Natl Acad Sci U S A. 2009 Aug 4;106(31):13004–9.

49. Fabra-García A, Yang AS, Behet MC, Yap Z, van Waardenburg Y, Kaviraj S, et al. Human antibodies against noncircumsporozoite proteins block Plasmodium falciparum parasite development in hepatocytes. JCI Insight. 2022 Mar 22;7(6):e153524.

50. McConville R, Krol JMM, Steel RWJ, O’Neill MT, Davey BK, Hodder AN, et al. Flp/FRT-mediated disruption of ptex150 and exp2 in Plasmodium falciparum sporozoites inhibits liver-stage development. Proc Natl Acad Sci U S A. 2024 July 9;121(28):e2403442121.

51. Aubrey BJ, Kelly GL, Kueh AJ, Brennan MS, O’Connor L, Milla L, et al. An inducible lentiviral guide RNA platform enables the identification of tumor-essential genes and tumor-promoting mutations in vivo. Cell Rep. 2015 Mar 3;10(8):1422–32.

52. Lupton EJ, Roth A, Patrapuvich R, Maher SP, Singh N, Sattabongkot J, et al. Enhancing longevity of Plasmodium vivax and P. falciparum sporozoites after dissection from mosquito salivary glands. Parasitol Int. 2015 Apr;64(2):211–8.

53. Roth A, Adapa SR, Zhang M, Liao X, Saxena V, Goffe R, et al. Unraveling the Plasmodium vivax sporozoite transcriptional journey from mosquito vector to human host. Sci Rep. 2018 Aug 15;8(1):12183.

54. Lord SJ, Velle KB, Mullins RD, Fritz-Laylin LK. SuperPlots: Communicating reproducibility and variability in cell biology. J Cell Biol. 2020 June 1;219(6):e202001064.

55. Torgler R, Bongfen SE, Romero JC, Tardivel A, Thome M, Corradin G. Sporozoite-mediated hepatocyte wounding limits Plasmodium parasite development via MyD88-mediated NF-kappa B activation and inducible NO synthase expression. J Immunol Baltim Md 1950. 2008 Mar 15;180(6):3990–9.

56. Liehl P, Zuzarte-Luís V, Chan J, Zillinger T, Baptista F, Carapau D, et al. Host-cell sensors for Plasmodium activate innate immunity against liver-stage infection. Nat Med. 2014 Jan;20(1):47–53.

57. Barata L, Houzé P, Boutbibe K, Zanghi G, Franetich JF, Mazier D, et al. In Vitro Analysis of the Interaction between Atovaquone and Proguanil against Liver Stage Malaria Parasites. Antimicrob Agents Chemother. 2016 July;60(7):4333–5.

58. Langlois AC, Marinach C, Manzoni G, Silvie O. Plasmodium sporozoites can invade hepatocytic cells independently of the Ephrin receptor A2. PloS One. 2018;13(7):e0200032.

59. Chainarin S, Jaihan U, Tapaopong P, Kongngen P, Kunkeaw N, Cui L, et al. Overexpression of hepatocyte EphA2 enhances liver-stage infection by Plasmodium vivax. Sci Rep. 2022 Dec 13;12(1):21542.

60. Xia M, Vago F, Han L, Huang P, Nguyen L, Boons GJ, et al. The αTSR Domain of Plasmodium Circumsporozoite Protein Bound Heparan Sulfates and Elicited High Titers of Sporozoite Binding Antibody After Displayed by Nanoparticles. Int J Nanomedicine. 2023;18:3087–107.

61. Coppi A, Tewari R, Bishop JR, Bennett BL, Lawrence R, Esko JD, et al. Heparan sulfate proteoglycans provide a signal to Plasmodium sporozoites to stop migrating and productively invade host cells. Cell Host Microbe. 2007 Nov 15;2(5):316–27.

62. Frevert U, Sinnis P, Cerami C, Shreffler W, Takacs B, Nussenzweig V. Malaria circumsporozoite protein binds to heparan sulfate proteoglycans associated with the surface membrane of hepatocytes. J Exp Med. 1993 May 1;177(5):1287–98.

63. Koike-Yusa H, Li Y, Tan EP, Velasco-Herrera MDC, Yusa K. Genome-wide recessive genetic screening in mammalian cells with a lentiviral CRISPR-guide RNA library. Nat Biotechnol. 2014 Mar;32(3):267–73.

64. Kueh AJ, Herold MJ. Using CRISPR/Cas9 Technology for Manipulating Cell Death Regulators. Methods Mol Biol Clifton NJ. 2016;1419:253–64.

65. Dull T, Zufferey R, Kelly M, Mandel RJ, Nguyen M, Trono D, et al. A third-generation lentivirus vector with a conditional packaging system. J Virol. 1998 Nov;72(11):8463–71.

66. Saliba KS, Jacobs-Lorena M. Production of Plasmodium falciparum gametocytes in vitro. Methods Mol Biol Clifton NJ. 2013;923:17–25.

67. Sinnis P, De La Vega P, Coppi A, Krzych U, Mota MM. Quantification of sporozoite invasion, migration, and development by microscopy and flow cytometry. Methods Mol Biol Clifton NJ. 2013;923:385–400.

68. Tewari K, Flynn BJ, Boscardin SB, Kastenmueller K, Salazar AM, Anderson CA, et al. Poly(I:C) is an effective adjuvant for antibody and multi-functional CD4+ T cell responses to Plasmodium falciparum circumsporozoite protein (CSP) and αDEC-CSP in non human primates. Vaccine. 2010 Oct 21;28(45):7256–66.

69. Austin LS, Kaushansky A, Kappe SHI. Susceptibility to Plasmodium liver stage infection is altered by hepatocyte polyploidy. Cell Microbiol. 2014 May;16(5):784–95.

70. Livingstone MC, Bitzer AA, Giri A, Luo K, Sankhala RS, Choe M, et al. In vitro and in vivo inhibition of malaria parasite infection by monoclonal antibodies against Plasmodium falciparum circumsporozoite protein (CSP). Sci Rep. 2021 Mar 5;11(1):5318.

71. Silvie O, Charrin S, Billard M, Franetich JF, Clark KL, van Gemert GJ, et al. Cholesterol contributes to the organization of tetraspanin-enriched microdomains and to CD81-dependent infection by malaria sporozoites. J Cell Sci. 2006 May 15;119(Pt 10):1992–2002.

72. Wilson DS, Hirosue S, Raczy MM, Bonilla-Ramirez L, Jeanbart L, Wang R, et al. Antigens reversibly conjugated to a polymeric glyco-adjuvant induce protective humoral and cellular immunity. Nat Mater. 2019 Feb;18(2):175–85.

73. van Schaijk BCL, Janse CJ, van Gemert GJ, van Dijk MR, Gego A, Franetich JF, et al. Gene disruption of Plasmodium falciparum p52 results in attenuation of malaria liver stage development in cultured primary human hepatocytes. PloS One. 2008;3(10):e3549.

74. Yang ASP, van Waardenburg YM, van de Vegte-Bolmer M, van Gemert GJA, Graumans W, de Wilt JHW, et al. Zonal human hepatocytes are differentially permissive to Plasmodium falciparum malaria parasites. EMBO J. 2021 Mar 15;40(6):e106583.

75. March S, Ng S, Velmurugan S, Galstian A, Shan J, Logan DJ, et al. A microscale human liver platform that supports the hepatic stages of Plasmodium falciparum and vivax. Cell Host Microbe. 2013 July 17;14(1):104–15.

76. March S, Nerurkar N, Jain A, Andrus L, Kim D, Whittaker CA, et al. Autonomous circadian rhythms in the human hepatocyte regulate hepatic drug metabolism and inflammatory responses. Sci Adv. 2024 Apr 26;10(17):eadm9281.

