## Supplementary Figure 1 for "Cas9-expressing HC-04 hepatocytes facilitate CRISPR-based analysis of *Plasmodium falciparum* sporozoite-host interactions"

**Contents**

Supplementary Figure 1

**HC-04 2B3** Δ***GALNT10***

```
DNA
sgRNA		--------------CCTTAC-CCCATGACCGATGCTGA----------
WT clone		GAGAACAAGGAAGACCTTAC-CCCATGACCGATGCTGAGAGAGTGGAT
KO clone allele 1	GAGAACAAGGAAGACCTTACCCCCATGACCGATGCTGAGAGAGTGGAT
KO clone allele 2	GAGAACAAGGAAGACCTTAC--CCATGACCGATGCTGAGAGAGTGGAT
Protein
sgRNA		----PYPMTDA---
WT clone		EQGRPYPMTDAERV
KO clone allele 1	EQGRPYPHDRC-ES
KO clone allele 2	EQGRPYP-PMLREW
```

**HC-04 2B3** Δ***CADM2***

```
DNA
sgRNA		-------------GAACTGCAATTTTGACCTGCAGG-------------
WT clone		GTTGTTGAAGGTGGAACTGCAATTTTGACCTGCAGGGTTGATCAAAATG
KO clone		GTTGTTGAAGGTGGAACTG-----------TGCAGGGTTGATCAAAATG
Protein
sgRNA		-----TAILTCR
WT clone		VVEGGTAILTCR
KO clone		VVEGGTVQG-SK
```

**HC-04 2B3** Δ***JAM2***

```
DNA
SgRNA		-----------------AGATGATAGATTTCAATATCCGG-----------------
WT clone		TTTTAAAAATCGAGCTGAGATGATAGATTTCAATATCCGGATCAAAAATGTGACAAG
KO clone		TTTTAAAAATCGAGCTGAGATGATAGATTTCA--ATCCGGATCAAAAATGTGACAAG
Protein
SgRNA		------MIDFNIR-----
WT clone		FKNRAEMIDFNIRIKNVT
KO clone		FKNRAEMIDFNPDQKCDK
```

**HC-04 2B3** Δ***POMT2***

```
DNA
sgRNA		---------------GGTCTTGCTGGCTACCTGAG-TGG----------------------
WT clone		CCTCAGATGCTGATAGGTCTTGCTGGCTACCTGAG-TGGATATGATGGTACCTTT------
KO clone		CCTCAGATGCTGATAGGTCTTGCTGGCTACCTTGAGTGGATATGATGGTACCTTTTTGTTC
Protein
sgRNA		-----GLAGYLS------
WT clone		PQMLIGLAGYLSGYDGTF
KO clone		PQMLIGLAGYLEWI-WYL
```

**HC-04 2B3** Δ***CLMP***

```
DNA
sgRNA		--------TGTCTACAATAACTTGACTGAGG-----------
WT clone		AGTCGTCATGTCTACAATAACTTGACTGAGGAACAGAAGGGC
KO clone		AGTCGTCATGTCTACAATTTCCTGGCAATTTCCTGGCAGGAG
Protein
sgRNA		---VYNNLTE---------------
WT clone		SRHVYNNLTEEQKGRVAFASNFLAG
KO clone		SRHVYNFLAISWQEMPPCRLNL-SP
```

**HC-04 2B3** Δ***DPY19L3***

```
DNA
sgRNA		------------ATCTCATTCAGAACAGAGTGTGG-----------------
WT clone		GTGGAGCGAGAAATCTCATTCAGAACAGAGTGTGGCCTGTATTACTCCTACT
KO clone		GTGGAGCGAGAAATCTCATTCAGAACAGT--GTGGCCTGTATTACTCCTACT
Protein
sgRNA		----ISFRTEC-----------
WT clone		VEREISFRTECGLYYSYYKQML
KO clone		VEREISFRTVWPVLLLLQADAA
```

**HC-04 2B3** Δ***KIRREL1***

```
DNA
sgRNA		--------AATTGCTGAAGGATGGGAAGAGG------------
WT clone		GTTCCAGGAATTGCTGAAGGATGGGAAGAGGGAGACCACCGTG
KO clone allele 1	GTTCCAGGAATTGCTGAAGGATGAAG--AGGGAGACCACCGTG
KO clone allele 2	GTTCCAGGAATTGCTGAAGGA--------GGGAGACCACCGTG
Protein
sgRNA		---------LLKDGKR-----
WT clone		SPTLVMFQELLKDGKRETTVS
KO clone allele 1	SPTLVMFQELLKDEEGDHREP
KO clone allele 2	SPTLVMFQELLKEGDHREPTA
```

**HC-04 2B3** Δ***DSG3***

```
DNA
sgRNA		------------CCAAGCAACCCAGAAAATCACCT----------
WT clone		TACTTCAGATTACCAAGCAACCCAGAAAATCACCTACCGAATCTC
KO clone		TACTTCAGATTACCC--------AGAAAATCACCTACCGAATCTC
Protein
sgRNA		-----QATQKIT---------------
WT clone		ITSDYQATQKITYRISGVGIDQPPFGI
KO clone		ITSDYPENHLPNLWSGNRSAAFWNLCC
```

**HC-04 2B3** Δ***PVRL2***

```
DNA
sgRNA		-------------CCTGAAGTGTCCATCTCCGGCTA------------------
WT clone		TCCCCCAGACCCTCCTGAAGTGTCCATCTCCGGCTAT-------------GATG
KO clone allele 1	TCCCCCAGACCCTCC-----TGTCCATCTCCGGCTAT-------------GATG
KO clone allele 2	TCCCCCAGACCCTCCTGGCTATGATGACAACTGGTACCTCGAGCAGCTGAAGC
Protein
sgRNA		-----PEVSISG-----
WT clone		FPPDPPEVSISGYDDNW
KO clone allele 1	FPPDPPVHLRL—QLVPR
KO clone allele 2	FPPDPPGYDDNWYLEQL
```

**HC-04 2B3** Δ***CLDN9***

```
DNA
sgRNA		---------------------CCTCCTGGTGGCCATCACAGGTG-------------------------
WT clone		CCTCCTGCTGGCCCTGCTTGGCCTCCTGGTGGCCATCACAGGTGCCCAGTGTACCACGTGTGTGGAGGA
KO clone allele 1	CCTCCTGCTGGCCCTGCTTG-----------GCCATCACAGGTGCCCAGTGTACCACGTGTGTGGAGGA
KO clone allele 2	CCTCCTGCTG----------------------------------GCCAGTGTACCACGTGTGTGGAGGA
Protein
sgRNA		-------LLVAITG---------
WT clone		LLLALLGLLVAITGAQCTTCVEDEGAKARIVLT
KO clone allele 1	LLLALLGHHRCPVYHVCGGRRCQGPYRAHRGGH
KO clone allele 2	LLLASVPRVWRTKVPRPVSCSPRGSSSSSPASW
```

**Supplementary Figure 1:** DNA and corresponding predicted amino acid sequences of human gene knockout cell lines generated from Cas9+ HC-04 2B3 parental cells by transduction, single cell cloning, expansion, and next generation sequencing of clonal cells.

Related to Figure 4.
